# Direct observation of adaptive tracking on ecological timescales in *Drosophila*

**DOI:** 10.1101/2021.04.27.441526

**Authors:** Seth M. Rudman, Sharon I. Greenblum, Subhash Rajpurohit, Nicolas J. Betancourt, Jinjoo Hanna, Susanne Tilk, Tuya Yokoyama, Dmitri A. Petrov, Paul Schmidt

## Abstract

Direct observation of evolution in response to natural environmental change can resolve fundamental questions about adaptation including its pace, temporal dynamics, and underlying phenotypic and genomic architecture. We tracked evolution of fitness-associated phenotypes and allele frequencies genome-wide in ten replicate field populations of *Drosophila melanogaster* over ten generations from summer to late fall. Adaptation was evident over each sampling interval (1-4 generations) with exceptionally rapid phenotypic adaptation and large allele frequency shifts at many independent loci. The direction and basis of the adaptive response shifted repeatedly over time, consistent with the action of strong and rapidly fluctuating selection. Overall, we find clear phenotypic and genomic evidence of adaptive tracking occurring contemporaneously with environmental change, demonstrating the temporally dynamic nature of adaptation.

**One sentence summary:** Rapid environmental change drives continuous phenotypic and polygenic adaptation, demonstrating the temporal dynamism of adaptation.

## Main text

Continuous adaptation in response to rapidly changing environmental conditions, termed adaptive tracking, could be a crucial mechanism by which populations respond to environmental change. Adaptive tracking has historically received little study due to the impression that adaptive evolutionary change is too slow to track complex and rapidly changing selection pressures in the wild (*1*). Moreover, theory suggests that variable and complex selective pressures should in general lead to the evolution of phenotypic plasticity or bet-hedging (*2, 3*). Yet, evidence of adaptation on ecological timescales from multiple longitudinal field studies and experiments demonstrates that adaptation can indeed occur very rapidly at individual traits or loci in response to strong environmental perturbations (*4–10*). Whether this translates into populations undergoing adaptive tracking in response to multifarious ecological changes, when theory predicts that pleiotropy should constrain natural selection and prevent adaptive tracking (*11, 12*), is unknown. If adaptive tracking does indeed occur in such situations, it would have broad implications for our understanding of the limits and pace of polygenic adaptation (*13*), the prevalence of fluctuating selection (*14*) and its role in the maintenance of genetic variation (*15*), and the importance of rapid adaptation in ecological outcomes (*16*).

To identify adaptive tracking it is necessary to directly measure phenotypic and genotypic evolution across replicate field populations in response to ongoing natural environmental change. Ideally an experimental system would provide: 1) the means for highly accurate measurements of even subtle, heritable shifts in key independent fitness-related phenotypes and loci under selection, 2) the ability to assay multiple replicate populations exhibiting some degree of ecological and environmental realism to detect parallel genetic and phenotypic changes indicative of adaptation (*17*), and 3) high resolution temporal sampling to quantify rapid fluctuations in the magnitude and direction of selection as environmental changes occur.

Here, we employ such an experimental system using field mesocosms to measure the extent, pace, repeatability, and genomic basis of adaptive tracking using *Drosophila melanogaster* in the naturally fluctuating, temperate environment of a single growing season in Pennsylvania, USA (*10, 18, 19*) (Fig. 1). The design precluded migration and allowed populations to expand to a large adult census size (on the order of 100,000 adults in each replicate at the maximum population size). To initiate the experiment, an outbred baseline population of *D. melanogaster* was derived from a set of 80 inbred strains originally collected in the spring from Pennsylvania (Table S1). Ten replicate cages were each founded on July 15th, 2014, with 1,000 individuals from the baseline population. All populations were tracked until the first hard frost on November 7th, 2014 and assayed at monthly intervals. Specifically, at four timepoints we measured the evolution of six complex, fitness-associated phenotypes, focusing on a set associated with either reproductive output or stress tolerance (Fig. 1). We repeatedly collected and reared individuals from each field cage in standard laboratory conditions (*i.e*., multi-generation common garden) to distinguish evolution from phenotypic plasticity and measured all phenotypes in the F3 generation. We also tracked changes in allele frequencies genome-wide in each replicate using pooled sequencing at five timepoints, employing haplotype-based allele frequency estimation (*20*) in order to generate highly accurate allele frequency trajectories.

**Figure 1:**
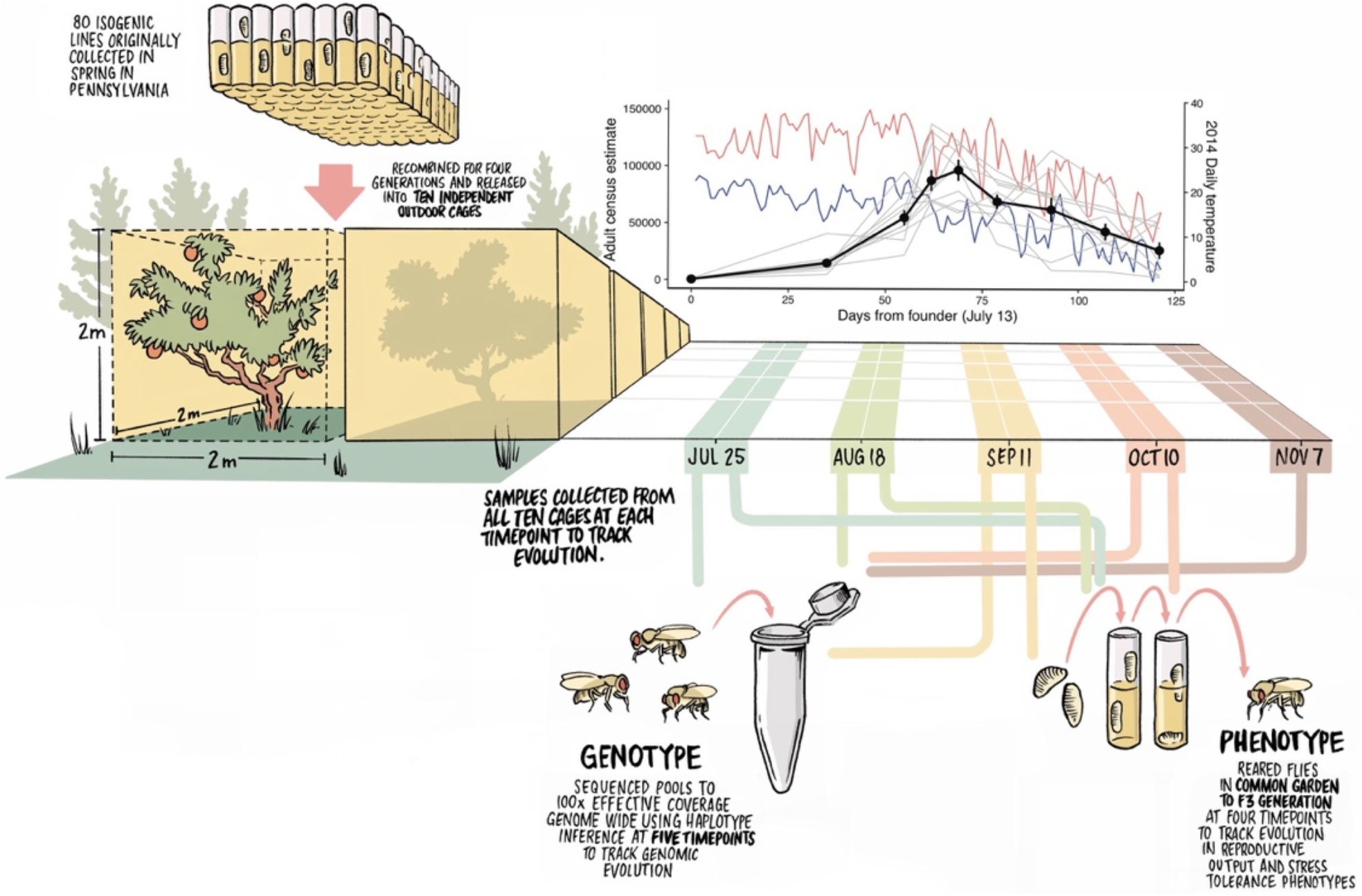
Experimental arena, design, and population dynamics. The experiment was designed to reflect ecological and evolutionary realism while testing for adaptation using replicate populations. 80 inbred lines originally collected in spring from an orchard in Pennsylvania were recombined and expanded for four generations into a genetically diverse outbred population in the laboratory. 500 males and 500 females from this outbred population were used to found 10 independent outdoor cages (2m x 2m x 2m). We measured daily minimum and maximum temperatures (blue and red lines, respectively) and estimated adult population size of each replicate over four months of seasonal change (mean: black line; per-replicate: gray lines). To study adaptation, we tracked phenotypic evolution by collecting eggs from each replicate, rearing them in common garden laboratory conditions for three generations, and then measuring six fitness-associated phenotypes. We conducted this procedure on the founder population and at four subsequent time points to measure phenotypic evolution over time. To study adaptation at the genomic level we sequenced pools of 100 females from each cage to > 100x effective coverage at five time points using haplotype inference [20] and assessed changes in allele frequencies.

Population dynamics were largely consistent among the replicates; population size increased sharply during summer, peaked in early fall, and then declined steadily as minimum daily temperatures declined in late fall (Fig. 1). These population dynamics mimic the patterns observed in *D. melanogaster* populations (*21*) and many other multivoltine organisms inhabiting temperate natural environments, with summer exponential growth, high densities in late summer to early fall, and late fall population declines. Egg production showed a similar pattern (Fig. S1), albeit at greater numbers, and overall recruitment from egg to adult was low (Fig. S2). Similarity in the ecological conditions among replicate populations, including abiotic factors (Fig. S3) and population dynamics (Fig. 1), suggests similar selective landscapes may have driven parallel evolution across replicates.

Phenotypic evolution was rapid and parallel, but temporal patterns varied across traits. In order to measure phenotypic evolution, we sampled individuals from the founding population and ~2,500 eggs from each cage at the first four time points (July 25, August 18, September 11, October 10), reared them in common garden laboratory condition for three generations, and assayed phenotypes in the F3 progeny (Fig. 1). For all six phenotypes, which are known to be polygenic and associated with fitness (*22*), we observed substantial trait evolution with an average of 23% change in the mean trait value for each cage across all phenotypes over each time interval. Variation in environmental parameters among cages did not implicate any individual factors as agents of selection (Fig. S4), perhaps due to the limited variation between cages or the complexity of the selective landscape. Prior experiments conducted in these mesocosms have found evidence of rapid adaptation in response to experimentally manipulated agents of divergent selection (*10, 19*).

All six phenotypes showed evidence of parallel evolution, indicative of adaptation, over time. Four of six phenotypes evolved rapidly, repeatedly, and in a consistent direction across the duration of the experiment (Fecundity: F_3,27_=43.75, p<0.0001; Egg size: F_3,27_=11.5, p<0.0001; Starvation: F_4,36_=129.05, p<0.0001; Chill coma recovery: F_4,36_=197.75, p<0.0001) (Fig. 2). The magnitude of change was often substantial: for example, the average increase in fecundity was 61% over each monthly sampling interval across replicates, representing 1-4 overlapping generations. Desiccation tolerance and development rate also evolved rapidly and in parallel (F_4,36_ =86.66, p<0.0001 Fig. 2C; F_4,36_=98.70, p<0.0001, Fig. 2F), but the direction of evolution varied over time. Fluctuations in the direction of evolution for these phenotypes had considerable effects on phenotypic trajectories; for desiccation tolerance the amount of evolution measured over the whole experiment (founder to October 10th) was less than what was observed over the first interval (founder to July 25th). Identifying the fitness effects of any specific instance of phenotypic evolution is complicated by underlying correlations among traits, pleiotropy, and an unknown and potentially temporally variable phenotype-to-fitness map but the pace and parallelism of phenotypic evolution is suggestive of strong links to fitness.

**Figure 2:**
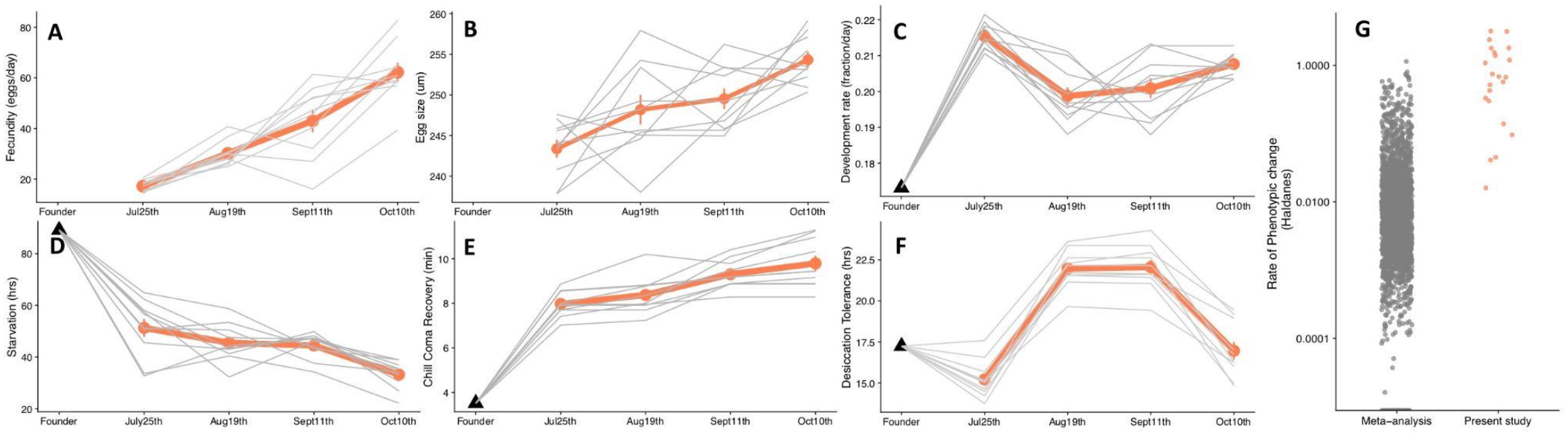
Parallel evolution of stress tolerance traits, reproductive output traits, and comparison of the rate of adaptation. Trajectories of phenotypic evolution for reproductive-associated (A, B, C) and stress resistance traits (D, E, F) as measured after three generations of common garden rearing. Panel A: mean fecundity as number of eggs/female/day, Panel B: mean egg size, Panel C: development rate as the fraction of development to pupation completed in one day (1/(total hours/24)). Panel D: starvation tolerance as time to death by starvation, Panel E: recovery time following chill coma, Panel F: desiccation tolerance as the time to death from desiccation. Black points are the mean phenotypes of the founding population, grey lines represent mean phenotypic trajectories of individual populations, and red lines are the mean of all cage means. Panel G: a comparison of the rates of adaptation from this experiment over individual intervals (red) to rates of phenotypic change from a prior meta-analysis (grey) [25].

The pace of parallel trait evolution observed over the short timescales examined in this study was unusually fast. As expected, we observed rapid parallel evolution when outbred laboratory populations were introduced into the field enclosures and adapted to the field environment (Founder → T_1_). However, we also observed evidence of rapid adaptation between intervals in the enclosures for all six phenotypes, with some showing reversals in the direction of evolution across intervals (Fig. 2 C&F). The rate of phenotypic adaptation, calculated in Haldanes (phenotypic evolution in units of standard deviations of the trait per generation (*23, 24*)), was computed as a mean change across replicates for each phenotype over each interval and across the whole experiment (Fig. 2G). The rate of adaptation over the whole experiment ranged from moderate to extremely fast for different traits (0 - 0.8 Haldanes) (*25*). However, when calculated over each sampling interval, the rate of adaptation was often comparable or faster than the fastest known pace of phenotypic change measured in any prior field study or experiment (Fig. 2G).

The pace, magnitude, and parallelism of the phenotypic evolution we observed is notable for three reasons. First, the evolutionary rates were calculated based on the phenotypic shifts of the F3 progeny in common garden conditions, thus excluding phenotypic plasticity as the driver of change. Second, because we focus only on the parallel phenotypic shifts across the cages, our estimates describe the rate of putatively adaptive phenotypic change. Third, these patterns of rapid adaptation were observed for multiple fitness-associated phenotypes, each with a complex and likely distinct genetic architecture (*26*).Overall, our results show that strong and temporally variable natural selection can consistently drive rapid and polygenic adaptation of multiple fitness associated phenotypes on the same timescale as the environmental change.

To investigate the genomic architecture underlying the observed rapid phenotypic adaptation, we performed whole-genome pooled sequencing of 100 randomly selected individuals from the baseline population and each replicate population at five timepoints across the experiment (Fig. 1). Allele frequencies at 1.9 M biallelic sites were inferred for each sample via haplotype inference using HAF-pipe [20] (Methods) at accuracy levels consistent with an ‘effective coverage’ of >100x (Supplementary Materials, Fig. S5, Table S2). This high-resolution dataset yielded strong evidence for rapid genome-wide evolution. Specifically, we observed that the genome-wide estimates of *F_ST_* between the founder population and all five monthly timepoints (mean 3.0 ± 0.2 x 10^-3^ standard error) exceeded expected margins of error based on technical and biological replicates (2.6 ± 0.24 x 10^-4^ and 1.8 ± 0.048 x 10^-3^ respectively, t-test p-values < 2×10^-8^, Fig. 3A). Furthermore, divergence from the founder population changed significantly over time both genome-wide (Kruskal-Wallace p-value for difference in means across timepoints: p < 2.3×10^-5^) and for individual chromosomes (p < 0.006, Fig. S6). Given the large population sizes (up to 10^5^) it is unlikely that such substantial evolutionary change can be attributed solely to random genetic drift.

**Figure 3:**
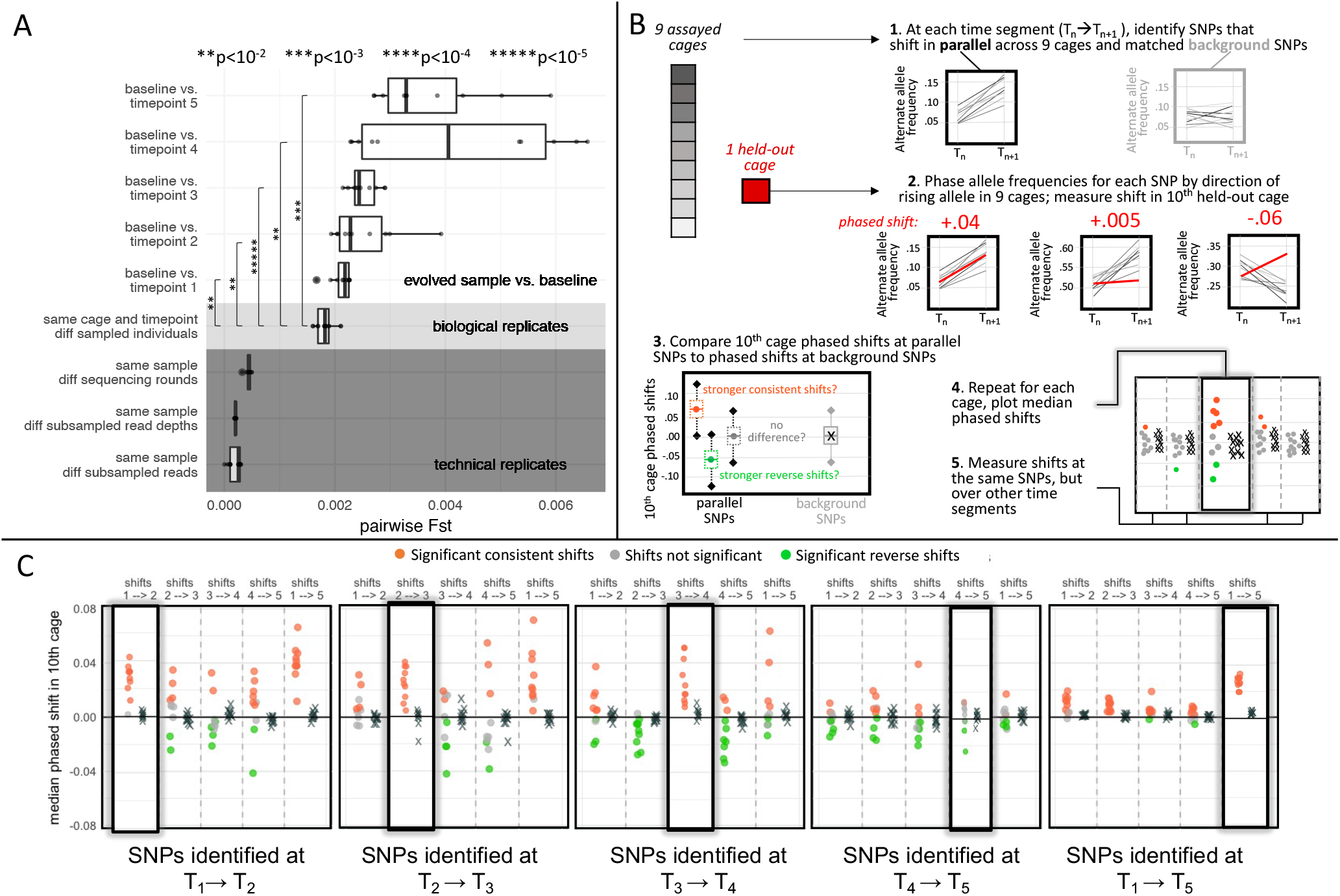
Using genomic data to test for evolutionary parallelism indicative of adaptation. A) Distributions of genome-wide mean pairwise Fst values between technical replicates (dark gray; same flies, different reads), biological replicates (light gray; different flies, same time point), and experimental samples from different timepoints compared to baseline (white). Note that negligible Fst values between pairs of technical replicates are consistent with extreme precision of HAFs, suggesting that the variance in allele frequency estimates for biological replicates is primarily driven by sampling of different individuals. Asterisks represent the significance of divergence over time compared to biological replicates (t-test). B) Graphical description of the leave-one out 10-fold cross-validation process for significant sites. In each round, significantly parallel sites (FDR <0.05, effect size>2%) at each time segment were identified using 9 of the 10 cages, then the shift at those sites in the 10th left-out cage was measured, after phasing such that positive values represent shifts in the same direction as the 9 assayed cages and negative values represent shifts in the reverse direction. The set of phased shifts at parallel sites was compared to phased shifts at background sites matched for chromosome and initial frequency and assigned to one of three significance bins: consistent (orange) or reverse (green), or no significant difference from background (gray). Shifts at these same sites over other time segments were also measured, phased, and assigned to significance bins. C) The median shift for each set of parallel sites (circles) and background sites (x marks) is plotted for each left-out cage. Each block of 5 panels represents shifts at the same sets of sites, those identified as parallel at the time segment labeled below the block. Shifts measured at that same time segment are highlighted in the panel with a dark shadowed outline.

Further examination of the magnitude and direction of evolution across the 10 replicate cages showed substantial genomic adaptation, as defined by parallel, and thus deterministic, allele frequency shifts across replicate cages. To test for parallel shifts, we used a leave-one-out cross validation approach. For each monthly time interval (T_i_ ➔ T_i+1_; i = 1,2,3,4), we used a generalized linear model (GLM) to identify sets of SNPs whose frequency shifted significantly across the 9 training cages, and then tested whether shifts at those SNPs in the 10^th^ left-out cage exceeded shifts at randomly-chosen matched control sites. Using this test, we found widespread parallel genomic adaptation for the first 3 sampling intervals (in 29 out of 30 leave one-out tests) (Fig.3C). The pattern of parallelism was muted and evolution was more idiosyncratic in T_4_➔ T_5_. We also repeated the procedure for SNPs that shifted across the whole experiment (T_1_ ➔ T_5_) and found a similarly strong signal of parallel adaptation (10 out of 10 tests). The magnitude of allele frequency shifts in each interval (2-8%) and over the whole experiment (2-5%) corresponds to very strong effective selection strength at the most parallel sites of ~10-50% per monthly interval (1-4 generations) (Materials and Methods). This pattern was largely repeated when analyzing sites from each chromosome individually (Fig. S7). In simulated populations with the same demographics as the experimental populations, allele frequency shifts of this magnitude were consistently achieved with selection coefficients <=50% on alleles spanning a wide range of initial frequencies over similar timescales (Supplementary Information; Table S3). The pronounced parallel shifts in allele frequency across independent populations demonstrate the strong action of natural selection.

Our cross-validation analysis also yielded clear evidence of variation in the magnitude and direction of selection over time, consistent with the observed patterns of phenotypic evolution for some traits (Fig. 2). Specifically, the leave-one-out analysis and the time series genomic data allowed us to examine the full trajectory of alleles detected at any specific time interval (T_det_). We found that these alleles do often shift significantly more than alleles at control sites (Fig 3C) at other time intervals; however, the nature of these shifts varied over time. In some left-out cages and at some time intervals, alleles shifted in a direction consistent with their behavior during T_det_ (orange points); however, in other cases the direction flipped, resulting in significant reverse shifts (green points). Reverse shifts were strongest for sites with T_det_ = T_3_→T_4_ (Aug→Sept) during the time when populations expanded most rapidly and reached their maximum. These T_3_→T_4_ parallel sites showed consistent shifts in the *opposite* direction during the preceding interval (T_2_→T_3_, July→Aug) when the populations were still expanding. In many cages, these sites also shifted in the opposite direction during the subsequent (T_4_→T_5_, Oct→Nov) interval when population sizes were declining. These patterns likely reflect the action of rapidly fluctuating selection over the 4 months of the experiment.

With a complex and rapidly fluctuating selective landscape adaptation occurs over multiple timescales simultaneously and inferred rates of adaptation are dependent on the timescale of sampling [13]. Our results clearly illustrate the extent to which lower-resolution temporal sampling would obscure the inference of adaptive tracking. While sites identified during individual time intervals often showed median shifts of >2% in a single month, the strongest parallel sites detected from lower-resolution sampling (i.e., sampling only at T_1_ and T_5_) showed smaller monotonic shifts at each interval (on average, 0.6% per month). Moreover, the magnitude of this discrepancy varied widely over time. Taken together, these results underscore the value of high-resolution temporal sampling in revealing the existence of both temporally variable and temporally consistent directional selective forces.

The number and genomic location of causal loci involved in adaptation is central to understanding the mechanics of the adaptive process [27]. To quantify the genomic architecture of adaptation we examined the distribution of parallel sites across the genome and developed an algorithm to differentiate putatively independent targets of selection from the sites whose shifts could largely be ascribed to linkage disequilibrium and genetic draft. We first fit allele frequencies from all 10 cages to a GLM and identified significantly parallel sites (Fig. S8) at each time segment (n=4,274) and across the whole experiment (n=5,036), yielding 9,310 significant shifts overall (Fig. 4A, Table S4; Materials and Methods). As expected from the leave one-out analysis, the sets were largely non-overlapping: the 9,310 detected parallel shifts occurred at 9,000 unique SNPs. Moreover, at each time interval and across the whole experiment, parallel sites were both strongly clustered (empirical p<0.01; Fig. S9) and showed significantly higher average linkage values than the matched control sites (paired t-test p-value < 10^-16^; Fig. S10) (Material and Methods), suggesting that most parallel sites were only linked to causal loci rather than being causal themselves.

**Figure 4:**
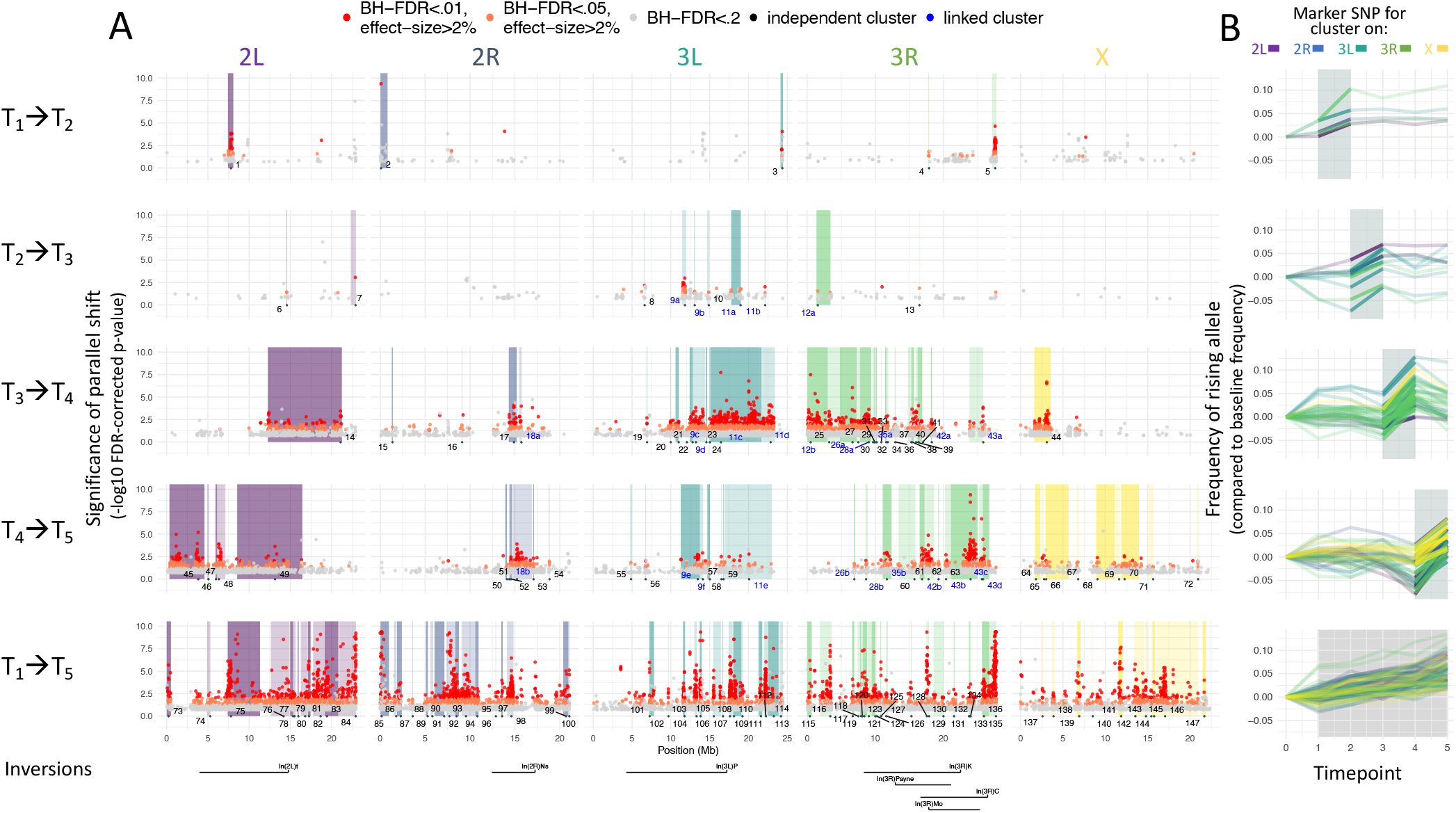
The genomic architectures of parallel allele frequency change over time. A) Manhattan plot of sites with significant parallel allele frequency shifts over time in 10 replicate cages. Each dot shows the -log10 of the FDR-corrected p-value (y-axis) corresponding to the significance of the allele frequency shift at a given SNP position (x-axis) over a given time segment of the experiment (rows). Only SNPs with an FDR <0.2 are shown, and dots are colored according to 3 significance bins (top legend). Shaded areas indicate regions of the genome that are likely driven by the same causal site, as defined by a clustering algorithm accounting for SNP linkage. Each clustered genome block is identified by a number marking the position of the top parallel SNP. Clusters from different time segments that are significantly linked (‘superclusters’) are given the same number, labeled in blue. The position of seven common chromosomal inversions are indicated below. B) Allele frequency trajectories are shown for the top marker SNP from each cluster. Each trajectory is translated to show allele frequency change relative to initial frequency in the baseline population, and phased to show the frequency of the rising allele at the time segment in which the cluster was identified. The time segment over which the SNPs were identified as outliers is shaded in gray.

We next identified the minimum number of independent genetic loci under selection using an algorithm that aggregated the parallel sites into clusters of linked sites (Materials and Methods, Fig. S10). This algorithm clustered 8,214 parallel SNPs detected across all the time segments (~90% of all SNPs at FDR <0.05) into 165 unlinked independent clusters (Fig. 4A, Table S5). These clusters were found on every chromosome and at every time segment, with an average of 4.5 clusters per chromosome per month. Simulations confirmed that while interference among multiple causal sites can temper shifts at any individual site, the number of clusters detected here still falls well within the realm of plausible selection landscapes. Specifically, when allele frequency trajectories for pairs, groups of 5, or groups of 10 selected loci were simulated simultaneously on the same chromosome, the majority (61.5%) of simulated selected sites required selection strengths no greater than s=0.5 to achieve a minimum shift of 2% per monthly time segment, and the vast majority (80.2%) required selection strengths no greater than s=1. Furthermore, although inversions can drive patterns of adaptation in *Drosophila* [28,29], no inversion markers were found among the parallel sites, and only 3 of the 165 clusters were strongly linked to inversions with average R^2^ > 0.1 (Table S7, Fig. S11). Combining clusters from all time segments, 61% of all assayed SNPs and 62% of the genome was contained in at least one cluster, highlighting the pervasive impact of short-term adaptive evolution at tens to hundreds of independent selected sites on allele frequencies genome-wide.

The genomic distribution and frequency shifts of these clusters suggested rapid changes in the targets and direction of selection over time. Specifically, 36 of the 90 clusters (40%) identified at a specific monthly time interval did not overlap any clusters identified at other monthly intervals, suggesting that selection at these loci was limited to one month. Among the remaining 54 clusters, only 27 (50%) contained SNPs that were significantly linked to SNPs in the cluster they overlapped. These 27 clusters formed 9 distinct ‘superclusters’ (Fig. 4) with high internal linkage, representing genomic regions in which allele frequencies shifted significantly in multiple monthly intervals. Strikingly, in 5 of the 6 superclusters involving a cluster from T_3_➔T_4_ linked to a cluster from T_4_➔T_5_, 90% of SNPs flipped direction between months, and in the 6th cluster >80% flipped direction, together totaling 10,464 SNPs that flipped direction (Fig. S12). A smaller majority of SNPs (67%) flipped in the supercluster formed by a cluster from T_2_➔T_3_ linked to a cluster from T_3_➔T_4_. Finally, in the two superclusters involving sets of linked clusters from 3 different time segments (T_2_➔T_3_, T_3_➔T_4_, T_4_➔T_5_), together covering over 5Mb of chromosome arm 3L, most SNPs (72% and 85%, respectively) flipped direction *twice*. We further confirmed that similar dynamics characterized the full set of putatively causal SNPs by choosing the SNP with the strongest parallelism p-value in each cluster and examining its trajectory (Fig. 4B). While the initial frequencies of these marker SNPs (Fig. S13) and exact shape of their trajectories varied widely, we observed a consistent trend: markers for the clusters identified at an individual monthly time interval often changed little during other months or even moved in the opposite direction (especially clusters identified at T_3_➔T_4_), whereas markers for clusters identified across the whole experiment tended to shift evenly and monotonically over time. The analysis of overlapping clusters and marker SNPs reveals similar patterns to individual SNP-based analyses, together supporting an oligogenic and rapid adaptive response to momentary selection pressures that often results in strong and rapidly fluctuating selection.

We next tested whether the identified genomic targets of this rapidly fluctuating selection are associated with any specific phenotypic traits or pathways. We specifically investigated the set of 111 genes - one per cluster - that overlapped with the cluster’s top marker SNP. This set of genes is strongly enriched (P < 0.001 in all cases) for genes with a known phenotypic effect (85 genes), and more specifically for genes involved in behavior (27 genes), cell-to-cell signaling (34 genes), neuronal function (25 genes) - and even more specifically, synaptic function (14 genes), and the CNS (21 genes) (Table S6). Many of these genes are crucial to core developmental and signaling pathways including the Wnt signaling pathway (genes *frizzled2* (the receptor of *wingless*), *armadillo* (beta-catenin), *sgg* (GSK3), *flo2* (long range Wnt signaling), *reck* (regulation of Wnt signaling), *huwe1* (negative regulation of Wnt signalling)), and dpp/BMP signaling (gene *tkv*). Strikingly, one cluster marker SNP is found in *SNF4Aγ*, the gamma subunit of the central metabolic switch kinase Adenosine 5’-monophosphate-activated protein kinase (AMPK). *SNF4Aγ* was one of two key genes found previously to be involved in adaptation to high temperature during experimental evolution of a sibling species, *D. simulans* (*30*). On balance these patterns suggest that the adaptive tracking in our outdoor mesocosms may be driven by the modulation of sensing and regulatory processes at the level of the nervous system, metabolism, and development that modify the way environmental cues are interpreted by the organism.

The phenotypic and genomic patterns observed in this study are consistent with a form of adaptive tracking in which (i) populations adapt in response to continuous environmental shifts, (ii) parallel evolution is driven by strong selection on multiple phenotypes and on a substantial number (tens to hundreds) of strongly selected genetic variants, (iii) the identity of the phenotypes and variants under selection changes considerably over short timescales, and (iv) selection operates at multiple timescales, acting in a consistent direction across the whole experiment on some variants and phenotypes, and rapidly fluctuating in direction and magnitude at others (*31*). This fluctuating selection leads to inferred rates of adaptation being slower when measured from the beginning to the end of the experiment as compared to single monthly intervals. The observed pattern that evolutionary rates are fastest when measured over shorter timescales may be driven by fluctuating selection (*13, 32*).

The pace, complex architecture of adaptation, and temporal evolution of particular phenotypes in our field cages are generally consistent with prior observations of seasonal evolution in natural temperate populations of *D. melanogaster* (*21, 33–35*). However, with additional temporal resolution and replication we detect rapidly fluctuating patterns of adaptation that suggest that populations of *D. melanogaster* are continuously and adaptively tracking the environment; this is surprising, but as we show not implausible given the timescale of environmental change (*36*). These patterns also imply that segregating functional variation is abundant and that much of the segregating variation in fitness is likely due to balancing selection (*37*), including temporally fluctuating selection that maintains genetic variation (*14, 38, 39*). The functional analysis of the genomic regions under selection further suggests that the rapid adaptation detected here is likely driven by modulation of high-level signaling pathways that feed into developmental and neuronal functions capable of modifying multiple phenotypes in a coordinated fashion. This may explain how selection can rapidly modify so many ostensibly unrelated phenotypes at the same time.

We show that it is possible to observe adaptive tracking in real time, providing a new lens to study the synchronous ecological and evolutionary dynamics of natural populations. We focus here on *D. melanogaster*, but the environmental and organismal features that gave rise to adaptive tracking, such as the presence of strongly shifting environmental pressures on generational time scales, are likely common (*7, 8, 40, 41*). Understanding the complex interplay among environmental change, population dynamics, standing genetic variation, and trait architecture that dictates the extent of adaptive tracking is a considerable challenge. Determining whether adaptive tracking is a general feature of natural populations and defining the factors that shape the extent of adaptive tracking could be transformative in understanding the generation and maintenance of biodiversity.

## Acknowledgements

We thank Andrew Berry, Moi Exposito-Alonso, Hunter Fraser, Dan Hartl, Jonathan Levine, Erin Mordecai, Dolph Schluter, and members of the Kelley and Cornejo labs, the King lab, the Petrov lab, the Schmidt lab, and two anonymous reviewers for helpful comments and discussions.

## Author Contributions

P.S. designed research. S.R., N.J.B., J.H. and P.S. conducted the experiment. T.Y and S.T. prepared sequencing libraries. S.M.R., S.I.G, D.A.P., and P.S. analyzed data, S.M.R, S.I.G, D.A.P, and P.S. wrote and revised the manuscript.

## Data and Code accessibility

Sequenced founder lines can be found at XXX. Sequencing data from evolved cages can be found at XXX. Scripts for the genomic analysis and simulations can be found at: https://github.com/greensii/dros-adaptive-tracking.

## SUPPLEMENTARY INFORMATION

### Materials and Methods

#### Establishment of experimental populations

To examine the pace, magnitude, parallelism, and genomic architecture of adaptation in response to a temporally variable environment we created a genetically diverse founder population that was seeded into each outdoor replicate. This outbred founder population was constructed from 80 fully sequenced *Drosophila melanogaster* inbred lines to facilitate the use of haplotype inference to attain high effective sequencing coverage. These inbred lines were derived from wild-caught individuals collected June 1, 2012 from Linvilla Orchards, Media PA USA (*1*). Each line was subsequently inbred for 20 generations by full-sib mating during which time they were maintained at 25 °C and fed ‘Spradling Cornmeal Recipe’ media. Then, 30-50 individuals from each line were pooled for whole genome sequencing. Sequencing and variant calling were performed as described in (*2*), with the addition that genomic DNA from certain lines was resequenced on an Illumina HiSeq X to increase coverage to a minimum of 10x for all lines. Mapped and de-duplicated bam files from all original and resequencing runs can be found on SRA under BioProject PRJNA722305 (Table S1). To initiate the baseline population in this experiment, we combined 10 males and 10 females from each of the 80 lines into large cages in May 2014. We allowed 4 generations of unmanipulated recombination and population expansion to facilitate recombination between lines before using 500 males and 500 female flies to found each of 10 field cages. Inbred lines have many deleterious alleles; purifying selection against deleterious alleles fixed during inbreeding was likely strong during lab outcrossing, and potentially, the early phase of the experiment.

Each field cage is a 2m x 2m x 2m mesh enclosure around a dwarf peach tree located outdoors (Philadelphia, PA) and features a natural insect and microbial community. The ground was fresh soil with clover planted as ground cover in each cage. The only food source and egg-laying substrate was 400ml of Drosophila media (‘Spradling cornmeal recipe’) contained in 900cm^3^ aluminum loaf pans that were added every second day for the duration of the experiment (July 13th - November 7th, 2014). Loaf pans of media within experimental cages were protected from rain and direct sun on shelving units oriented away from direct sunlight.

#### Measurement of population size and evolution of fitness associated phenotypes

Census size of adults was estimated in each replicate over the course of the experiment by photographing an equal amount of the surface area (approximately 2.5%) of the ceiling in each cage at dusk (12 total census estimates per cage). The number of adult *D. melanogaster* in each of 8 standardized photographs in each estimate for each cage was counted and multiplied by 40 to correct for total surface area and obtain census estimates. Egg production was estimated every second day by counting the eggs present on a 1/24^th^ portion of the exposed surface of the media.

To assess the rate and direction of phenotypic evolution over the course of the experiment we collected ~2500 eggs from each cage, brought them to the laboratory, and reared them for an additional 2 generations in a common garden (25°C, 12L:12D) while maintaining population sizes at ~2500 individuals. Fitness-associated phenotypes were measured on density and age-controlled replicates in the F3 generation. Fecundity was measured as the total number of eggs laid by a group of five females, counted each day for a period of three days, with twenty replicate vials for each cage at each time. Egg length was measured using a microscope and image processing software (*3*) on at least 15 eggs (average of 27) from each cage at each time point. Larval development rate was tracked as the time from when eggs were laid until pupation in three replicate vials from each cage at each time point with 30 eggs in each vial. Starvation tolerance was measured as time to starvation in three replicate vials containing moist cotton (1.5 ml water) (following (*4*)) and 10 female flies with three replicates for each cage at each time point. Desiccation tolerance was measured as time to death in desiccation chambers containing 10 female flies with three replicates for each cage at each time point (*4*). Chill coma recovery was measured as the time it took for flies buried in ice and placed in a 4°C incubator for 2h to resume an upright stance at 25°C (1). This was measured using groups of 10 female flies for each cage at each time point that had been allowed at least 24hrs to recover from light CO_2_ anesthetic. We also attempted to measure evolution in heat knockdown. However, the assay temperature we used for the founder population, a stressor that caused 50% of flies to knockdown by 12 minutes, was not sufficiently hot to cause knockdown by the second sample period. Thus, although we cannot quantify it, heat tolerance evolved rapidly. We assayed each of the remaining phenotypes in the founding population (founder assays failed for fecundity and egg size) and at four times during the experiment: day 11 (7/25/14), day 38 (8/19/14), day 61 (9/11/14), and day 90 (10/10/14). Census and phenotypic evolution data have been uploaded to Dryad.

#### Calculation of evolutionary rates and statistical analysis of phenotypic data to test for evolutionary parallelism

We calculated evolutionary rates in Haldanes by dividing the trait change over each interval by the pooled standard deviation and then by the number of generations elapsed (*5,6*). We calculated the rate of adaptation as the parallel change across replicates. To do so we took the average trait change across all 10 replicates and calculated a single rate in Haldanes. Haldanes were calculated for all six phenotypes for each experimental interval and over the whole experiment. We compared the rates of evolution measured in our experiment to those from a meta-analysis of evolutionary rates from field populations that focused on contemporary evolution (less than 200 generations) (*7*). The meta-analysis was focused on phenotypic change, which includes both genetic and environmental (plastic) effects, as few prior studies used common garden experiments to measure the rate of evolution.

To test for parallel phenotypic evolution in each of the six phenotypes we carried out separate linear mixed effect models (e.g. lme(phenotype measured ~ time, random=~1|cage/time)) and tested for significance using an anova (nlme and R respectively).

#### Genomic sequencing, SNP calls, and bioinformatic analysis

100 female flies from each of the 10 field cages were sampled at 5 monthly time points. Individuals from each sample were pooled and libraries were prepped using a Covaris protocol, then size-selected using an e-gel. Two e-gel bands from each sample were sequenced separately (1 from the 450-500 band and 1 from the 500-550 band) on a HiSeq3000 with 150-bp paired-end reads. Truseq adapter sequences and bases with quality <20 were trimmed with skewer (*8*) and overlapping forward and reverse reads were merged using PEAR (*9*). Resulting reads were mapped to the *Drosophila melanogaster* reference genome v5.39 with BWA (default parameters) (*10*). Reads were deduplicated using Picardtools and realigned around indels using GATK v4 (*11*). Pairs of bam files from the same sample were merged with samtools (*12*). Final average per-sample read depth was 7.3x +/- standard deviation of 2.0x. Haplotype-derived allele frequencies (HAFs) were then calculated via local inference with HAF-pipe (*2*) using the 80 genotyped founder strains. Haplotype inference was conducted in sliding windows across the genome, using the adaptive window size option in HAF-pipe to reflect the expected length of un-recombined haplotype blocks given the estimated number of generations since population founding. Heterozygous calls in the founder lines were included in the inference calculation, and missing calls were imputed using HAF-pipe’s ‘npute’ option. HAFs from all samples were compiled and filtered to contain only sites at which at least one baseline sample and at least one evolved cage sample had a minor allele frequency >1%.

#### High coverage sequencing

4 biological replicate samples from the baseline population, each a random sample of 100 flies from the same baseline population, were sequenced at high coverage. Baseline library preps were created using a modified Nextera protocol (*11*) and sequenced on a HiSeq4000 with target 100x coverage. Additionally, timepoint-5 evolved samples from 8 of the 10 cages were re-sequenced at high coverage (in addition to separate sequencing at low coverage with the rest of the evolved samples) using a KAPA hyperprep and a HighSeq4000. Processing for both the baseline and high-coverage timepoint-5 samples followed the same workflow. All adapter sequences were trimmed with skewer (*7*) with default parameters and minimum quality Q=20. Overlapping forward and reverse reads were merged using PEAR (*8*). Resulting reads were mapped to the *Drosophila melanogaster* reference genome v5.39 with BWA (default parameters). Reads were deduplicated using Picardtools and realigned around indels using GATK v4. Raw allele frequencies at each SNP site were then calculated using Popoolation (*12*) and custom bash scripts.

#### Analysis of HAF accuracy

Our approach relies on a previously published expectation-maximization algorithm for inferring the frequency of individual founder haplotype blocks in each pooled sample, which we then translate to population allele frequency estimates using weighted sums of founder genotypes. This approach was described in detail in (*2*), where we demonstrated via simulations that HAFs calculated from read depths ~5x can be as accurate as raw allele frequencies calculated from read depths >100x, and that high accuracy is maintained for >50 generations in *Drosophila*, although recombination does impact accuracy over time. As our experiment lasted only 10-15 generations, we expected that this approach would yield reliable allele frequencies suitable for downstream analysis. In fact, using the predictive tool described in (*2*) (https://ec-calculator.shinyapps.io/shinyapp/) to predict the expected ‘effective coverage’ of our HAFs from experimental parameters, accuracy estimates ranged from 106x for the most shallowly sequenced sample on the X chromosome (where SNP density is lowest, leaving fewer discriminatory sites for haplotype inference) to 369x for the deepest sequenced sample on chromosome 3R (Table S2).

However, to validate that HAF accuracy was sufficiently high with empirical (rather than simulated) data, and to confirm that this approach does not lead to biased estimates as recombination progresses, we re-sequenced 8 of the timepoint-5 samples at high-coverage and compared allele frequencies calculated from raw reads (‘raw AFs’; i.e., calculated from the proportion of alternate alleles at each site, without haplotype inference) to HAFs calculated from the same samples. Importantly the raw AFs and HAFs were calculated from distinct sets of reads (different aliquots of genomic DNA from the same individuals), and were thus independent estimates. Furthermore, while neither HAFs nor raw AFs represent ground truth allele frequencies for the sampled individuals, they each contain different sources of error. Thus, we would expect that the accuracy of HAFs would be reflected in a strong correlation with raw AFs at the highest read depths, since they are both faithful representations of the same signal, while if HAFs were systematically biased, increasing the raw AF read depth would not improve the correlation. To test this, sites in all 8 samples were binned by their read depth in the high coverage version of each sample, and then 50,000 sites were sampled randomly from each bin across all samples. Fig S1A shows density heatmaps of allele frequencies vs HAFs calculated at the same site in the same sample for sites in 4 different read depth bins. We observed that as raw read depth increased, raw allele frequencies more closely matched HAFs, as apparent from lower variance around the diagonal in the heatmap and a stronger correlation coefficient. To further confirm that there was no systematic bias in HAFs compared to raw allele frequencies, we plotted the smoothed line of best fit (using the function geom_smooth from the ggplot2 R package) separately for each read depth bin (Fig S1B). Indeed, for sites in the highest read depth bin, the line of best fit is almost exactly on the diagonal. Since our analysis relies not just on estimating allele frequencies correctly, but on detecting subtle shifts in allele frequency over time, we generated the same set of plots and correlations for the shift between baseline and timepoint 5 calculated from raw AFs vs HAFs (Fig. S1C-D). We observed the same pattern, in which concordance between raw AFs and HAFs improved with higher raw AF read depth, though the correlation coefficients overall were not as strong. These reduced correlation coefficients are expected given that the vast majority of shifts are very small and the dynamic range of values is reduced. Nevertheless, the consistent increase in correlation coefficient across read depth bins is consistent with HAF accuracy reaching effective coverages >115x (the highest read depths observed in the raw AFs). Finally, to assess the fine-scale resolution of HAFs, sites with raw read depth >115x and shifts <=10% were binned by raw AF shift to the nearest 1%, and boxplots were generated of HAF shifts at the sites in each bin (Fig S1E). The means of the HAF shifts in each bin rose significantly across each consecutive bin (all t-test p-values <.05), suggesting that HAFs provide the resolution necessary to distinguish shifts that differ by ~1%.

#### Identifying significant parallel SNPs

A generalized linear model (GLM) with a quasibinomial error model was fit to allele frequencies at each SNP to assess the parallelism of shifts in allele frequency across cages over each time interval. To account for sampling of chromosomes, all allele frequencies were first scaled and rounded to counts out of N_effective_, where *n* is the number of individuals sampled from the population (100 for all samples), *rd* is the true read depth, and N_effective_ = 2*n*\**rd*−1 / 2*n*+*rd*. A site was considered significantly parallel if it showed 1) at least 2% average change in allele frequency over the time interval and 2) Benjamini-Hochberg false discovery rate corrected p-value <.05 from the GLM test of parallelism. We also created an empirical false discovery rate correction by shuffling the sample time point labels and re-running GLMs, however this rate proved to be less stringent and therefore was not used in the analysis.

#### Leave-one out cross validation analysis

In each round, a GLM was fit using allele frequencies from 9 training cages, and parallel sites were identified at each time segment as described above. For each parallel site, a matched control site was identified on the same chromosome that had an initial frequency in the baseline population within 5% of the parallel site. At each parallel and control site, the allele frequency shift over each time segment in the 10th left-out cage was calculated and phased such that a shift in the same direction as the training cages was given a positive sign and a shift in the opposite direction was given a negative sign. A t-test was conducted for each time segment to determine if the set of phased shifts at parallel sites was significantly different than shifts across all control sites. In Figure 3, we plotted the median phased shift for each set of sites at each time segment, and colored the point for parallel sites if the t-test p-value was < 0.05 after false discovery rate correction.

#### Forward simulation of selection in replicate populations

Simulations of allele frequency dynamics associated with rapid adaptation were performed with the software tool forqs (*13*), which simulates recombination of haplotype chunks in the presence of zero or more selected alleles in a randomly mating population over a specified number of non-overlapping generations. We first chose a set of 100 sites from across the allele frequency spectrum on which to focus our simulations. To do so, we divided all segregating sites in the experimental founder population into 100 equidistant bins according to their alternate allele frequency across the 80 founder lines, and then randomly selected 1 site from each bin. Then, separately for each site, we used forqs to simulate allele frequency trajectories from 10 independent populations of 100,000 individuals over 3 generations of neutral ‘burn-in’ and 4 generations of constant directed selection on one of these 100 sites. In each simulation, the 100,000 individuals in each of the 10 populations were each assigned to carry the alleles of a randomly selected homozygous founder strain, which were supplied to forqs via an ms file. Simulations for each site were repeated with a range of selection coefficients between *s*=0.05 and *s*=1, in which homozygous reference, heterozygous, and homozygous alternate genotypes were assigned a selective advantage equal to 1, 1+*s/2*, or 1+*s* respectively. In each simulation we also tracked the frequency of neutral (ie *s*=0) marker sites located approximately 5kb away from each selected site. Environmental variance between populations was set to 0.05. To be conservative, in our simulations we referred to the female *D. melanogaster* recombination rate map (*14*) for all individuals, and simulated truncation selection in which the top 25% of individuals contribute to the next generation. After simulating selection on each site individually, we then randomly grouped the sites into pairs, sets of 5 sites, and sets of 10 sites, and repeated the simulations with multiple sites under selection with the same strength, each contributing independently to a single additive trait. After simulation for each site or set of sites, allele frequencies at each selected site and each marker site were averaged across the 10 replicate populations and the minimum selection coefficient was identified at which average allele frequency shifted by at least 2% over the course of the 4 generations of selection. Results are presented in Table S3.

#### Defining SNP clusters

A GLM model was fit to allele frequencies from all 10 cages at each site as described above, to assess the parallelism of the shift over each time interval. Each site was assigned a score for each time interval according to the following criteria: 0 = [FDR >0.2], 1 = [FDR<.2 or FDR>.2 and effect size <2%], 2 = [FDR<.05, effect size >2%], 3 = [FDR<.01, effect size >2%]. While only sites receiving a score of 2 or 3 were defined as ‘significant’ in the analysis, lower scoring sites were helpful in identifying large regions of elevated parallelism. Average SNP scores were calculated for sliding windows of 500 SNPs (offset=100 SNPs), and significantly enriched windows were defined as those with an empirical FDR <.05 compared to the distribution of window scores obtained by randomly shuffling sites across the genome. Overlapping enriched windows were then merged. Next, linkage was calculated between all pairs of significant SNPs less than 3 Mb apart from the same time interval. Linkage was defined as the squared correlation coefficient from a Pearson correlation of founder genotypes at the two sites, with genotypes coded as 0, 0.5, 1, or NA for missing data. Neighboring windows with average SNP-pair linkage >0.03 were merged into clusters, and the process was repeated iteratively until no neighboring clusters within 3Mb exceeded an average linkage of 0.03.

#### Defining superclusters

A list was generated of all pairs of clusters identified at different time segments that overlapped by at least one SNP. Clusters identified across the whole experiment (T_1_→T_5_) were excluded from this list, resulting in 44 pairs of overlapping clusters. For each pair of clusters, linkage (R^2^) values between all inter-cluster pairs of significant SNPs within 3Mb of each other were calculated and compared to linkage values for a set of randomly selected control SNP pairs matched for chromosome, initial frequencies, and inter-SNP distance. If linkage values for the cluster SNPs were significantly higher than linkage values for the matched control SNPs (Benjamini-Hochsburg FDR-corrected t-test p-value <.05), the clusters were considered significantly linked. Any individual pairs of linked clusters that shared a cluster in common were merged into linked cluster sets to form the final list of superclusters.

#### Assessing the influence of inversions

Inversion markers (*15*) were used to assess the linkage of each cluster to each inversion on the same chromosome. Markers were filtered to SNPs segregating in our baseline population. Because subsets of markers for the same inversion often showed disparate allele frequency trajectories in our data (and thus may not be reliable markers of the inversion among the inbred lines used to found our population), we filtered markers for each inversion to those that showed strong linkage (R^2^> 0.5) to at least half of the other markers for that inversion (see Table S7 for inversion marker counts before and after filtering). We then calculated the linkage between all significantly parallel SNPs and any inversion markers up to 3Mb away.

### Supplementary Tables

Table S1. List of inbred *Drosophila melanogaster* lines and SRA accession numbers used in this study.

(*see excel spreadsheet*)

**Table S2.**
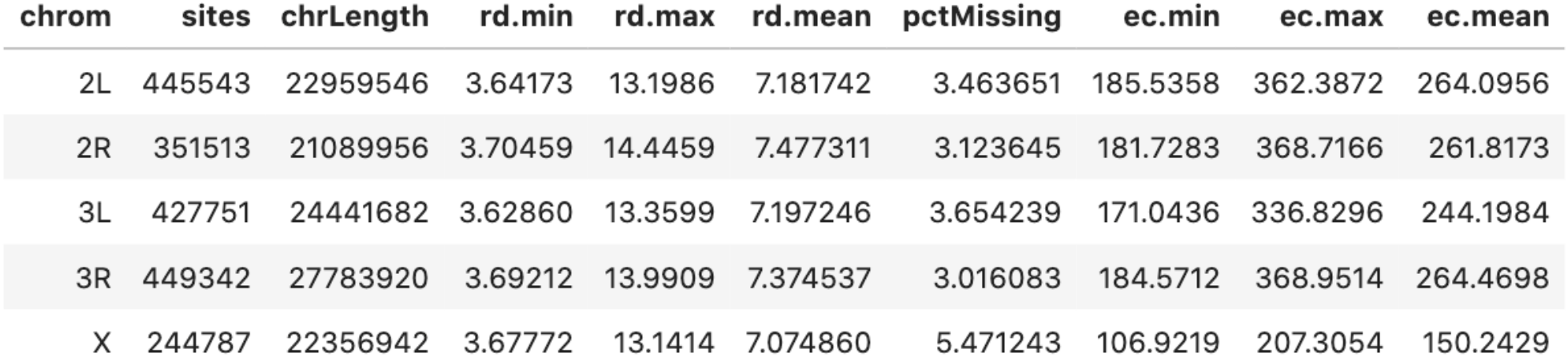
Predicted effective coverage (‘ec’) calculated from the density of sites per chromosome, percent of founder genotype calls that were missing, and the minimum, maximum, and mean chromosome-wide average read depth across samples according to the model described in (Tilk et al. 2019).

Table S3. Results of forward simulations of selection over 4 generations. For each selected site (left side) or marker site 5kb away (right side), the table lists the minimum selection coefficient required to shift allele frequency by 2% when the selected site was the only site under selection (first column), or was part of a multi-site selection regime (subsequent columns). NA indicates that no selection strengths tested resulted in a sufficient parallel shift.

(*see excel spreadsheet*)

**Table S4.** Counts of sites with significant (FDR<.05, effect size >2%) parallel allele frequency shift across 10 replicate cages at each time segment, on each chromosome.

| **Sig Site Count** | **2L** | **2R** | **3L** | **3R** | **X** | **Total** |
| --- | --- | --- | --- | --- | --- | --- |
| Timepoint 1 --> 2 | 42 | 4 | 6 | 67 | 4 | 123 |
| Timepoint 2 --> 3 | 4 | 0 | 109 | 5 | 0 | 118 |
| Timepoint 3 --> 4 | 178 | 109 | 1453 | 858 | 137 | 2735 |
| Timepoint 4 --> 5 | 306 | 156 | 93 | 642 | 101 | 1298 |
| Timepoint 1 --> 5 | 1294 | 1085 | 997 | 1146 | 514 | 5036 |
| Total | 1824 | 1354 | 2658 | 2718 | 756 | 9310 |
| Unique sites | 1783 | 1326 | 2541 | 2602 | 748 | 9000 |
| % of All sites | 0.4% | 0.38% | 0.59% | 0.58% | 0.31% | 0.47% |

**Table S5.** Counts of clusters identified at each time segment on each chromosome.

| **Cluster Count** | **2L** | **2R** | **3L** | **3R** | **X** | **All** |
| --- | --- | --- | --- | --- | --- | --- |
| Timepoint 1 --> 2 | 1 | 1 | 1 | 2 | 0 | 5 |
| Timepoint 2 --> 3 | 2 | 0 | 6 | 2 | 0 | 10 |
| Timepoint 3 --> 4 | 1 | 4 | 10 | 20 | 1 | 36 |
| Timepoint 4 --> 5 | 5 | 6 | 8 | 11 | 9 | 39 |
| Timepoint 1 --> 5 | 12 | 16 | 14 | 22 | 11 | 75 |
| All | 21 | 27 | 39 | 57 | 21 | 165 |

Table S6. Gene associations and annotations for the single marker SNP in each cluster with the strongest parallelism score. Columns marked with an asterisk represent phenotypic associations of marker genes obtained from http://evol.nhri.org.tw/phenome2/ (Weng et al 2017).

(*see excel spreadsheet*)

**Table S7.** Table of inversion marker counts. Segregating markers could be detected as bi-allelic SNPs in the baseline population, while filtered markers showed high correlation (R^2^>0.5) with each other across all sampled cages during the course of the experiment.

| Inversion | All Markers | Segregating Markers | Filtered Markers |
| --- | --- | --- | --- |
| In(2L)t | 16 | 16 | 16 |
| In(2R)Ns | 67 | 22 | 19 |
| In(3R)C | 144 | 10 | 0 |
| In(3R)K | 4 | 1 | 1 |
| In(3R)Mo | 150 | 73 | 64 |
| In(3R)Payne | 19 | 12 | 11 |

### Supplementary Figures

**Fig. S1:**
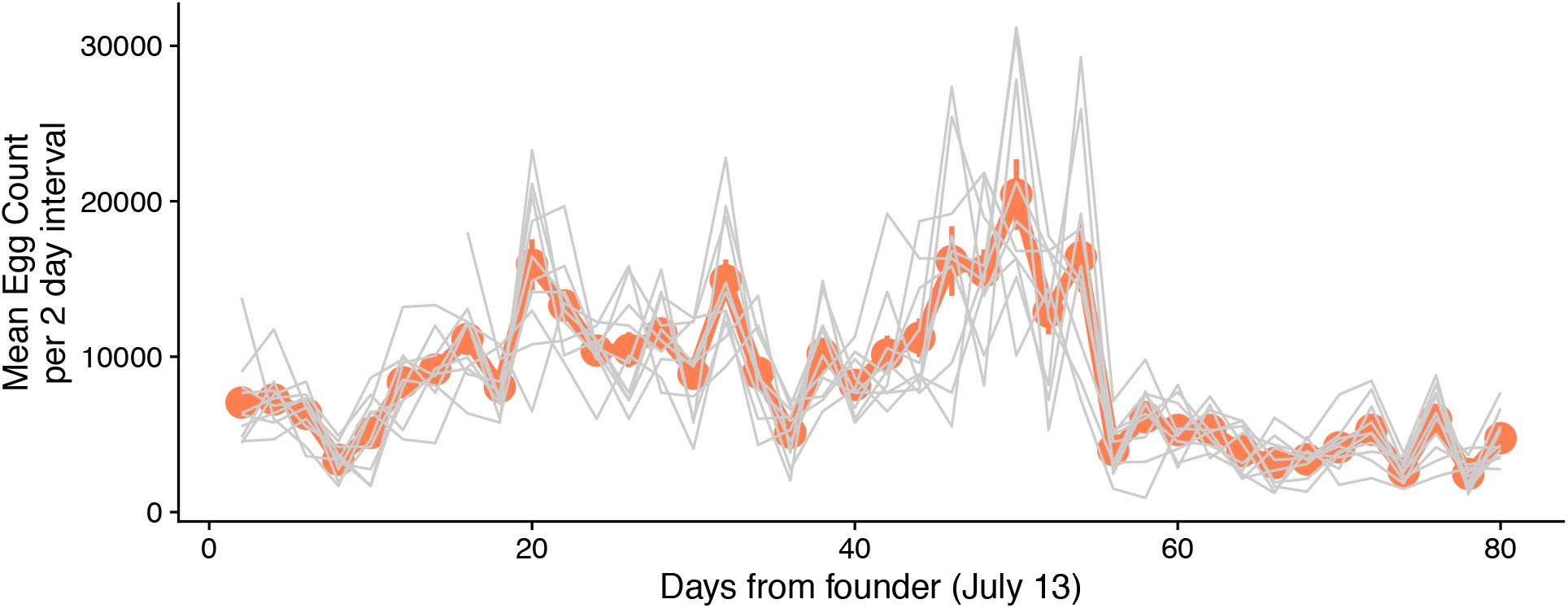
Eggs were estimated by counting the number on 1/24^th^ of the food loaf pan every second day during the experiment. Plotted here are the means (orange line) and individual cage value for egg production for each 2 day period.

**Figure S2:**
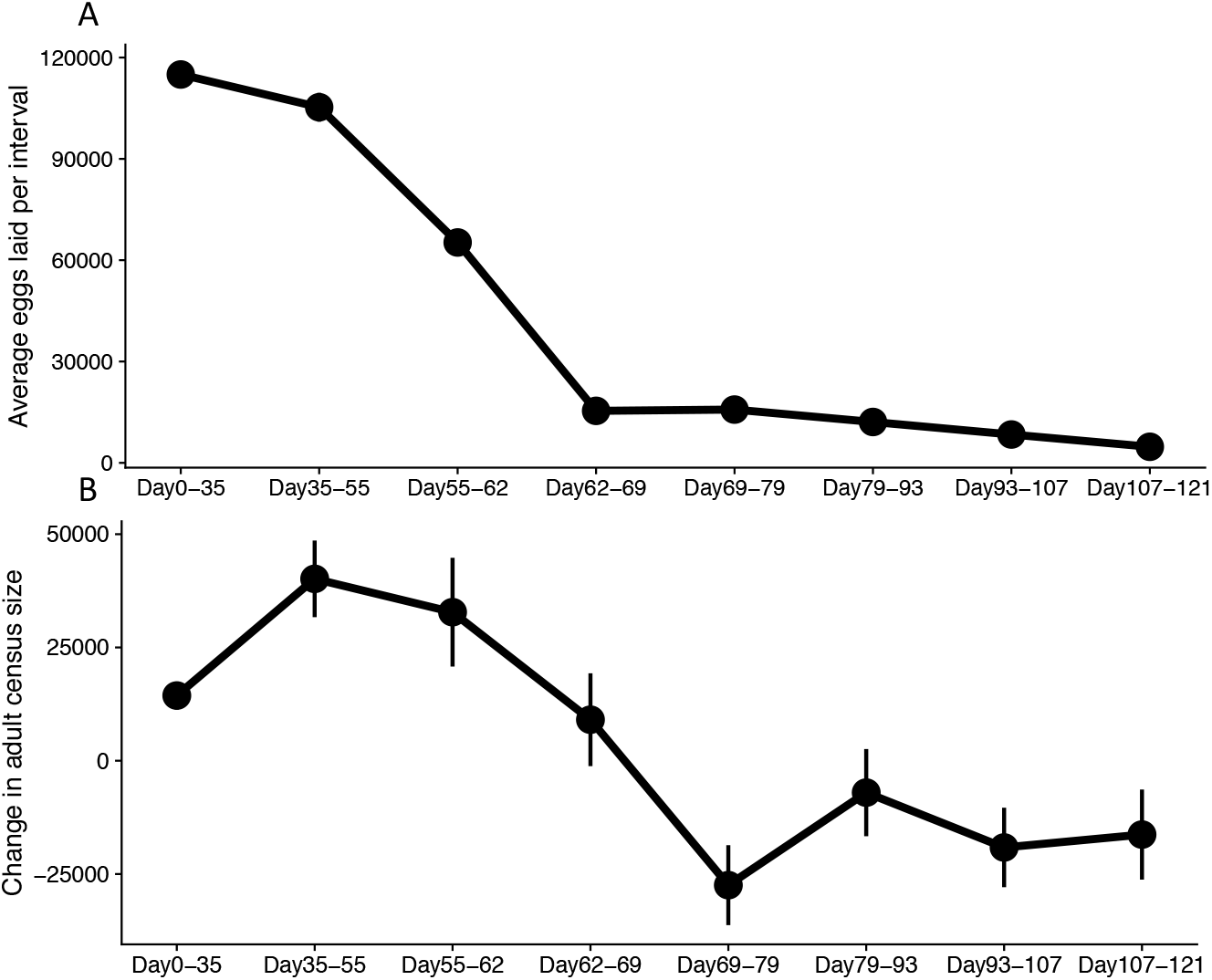
To visualize recruitment from egg to adult we have plotted: A) The total number of eggs that could have matured to adulthood between each adult census estimate B) The change in adult population size between each census estimate. For both A and B means with standard errors are plotted.

**Figure S3:**
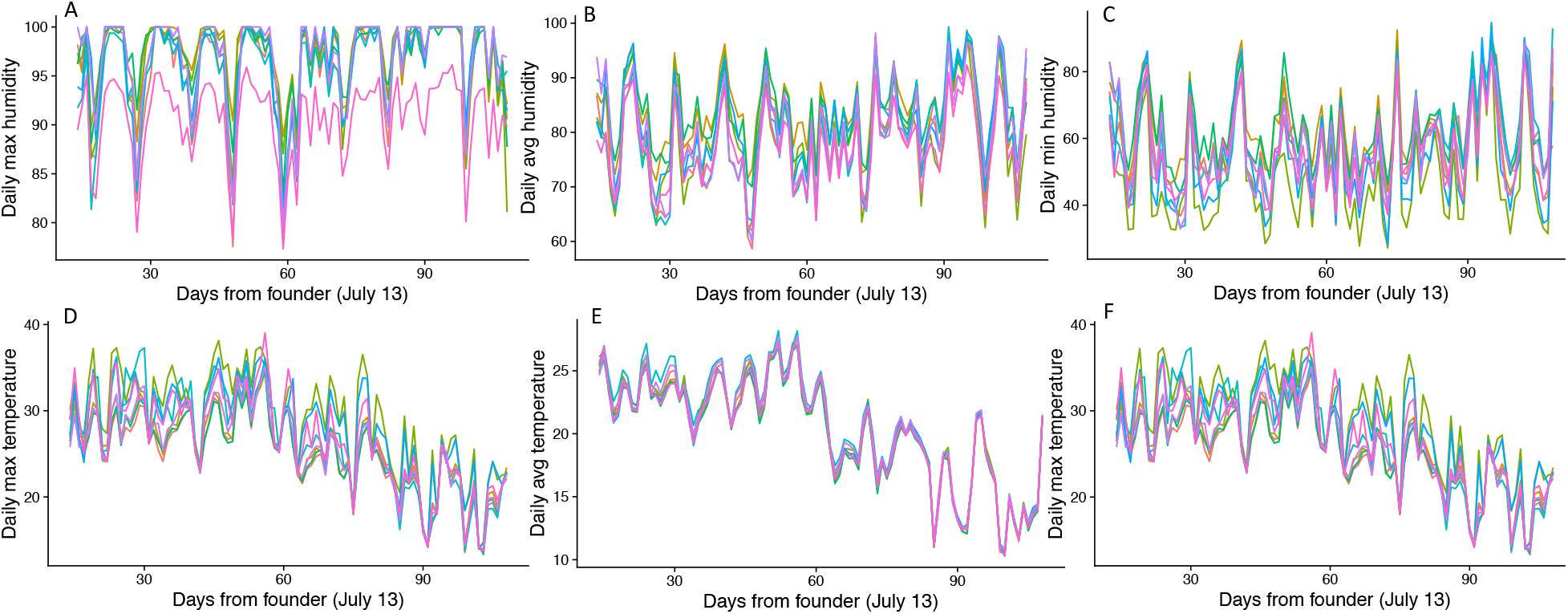
Panels A-C show cage by cage variation in daily relative humidity (A=maximum, B=average, C=minimum). Panels D-F show cage by cage variation in daily temperature (D=maximum, E=average, F=minimum). Temperature and humidity loggers in 8 of 10 cages collected complete data and are included here. Cage level variation is modest overall, maintaining the expectation that independent replicate populations may show parallel evolutionary trajectories.

**Figure S4:**
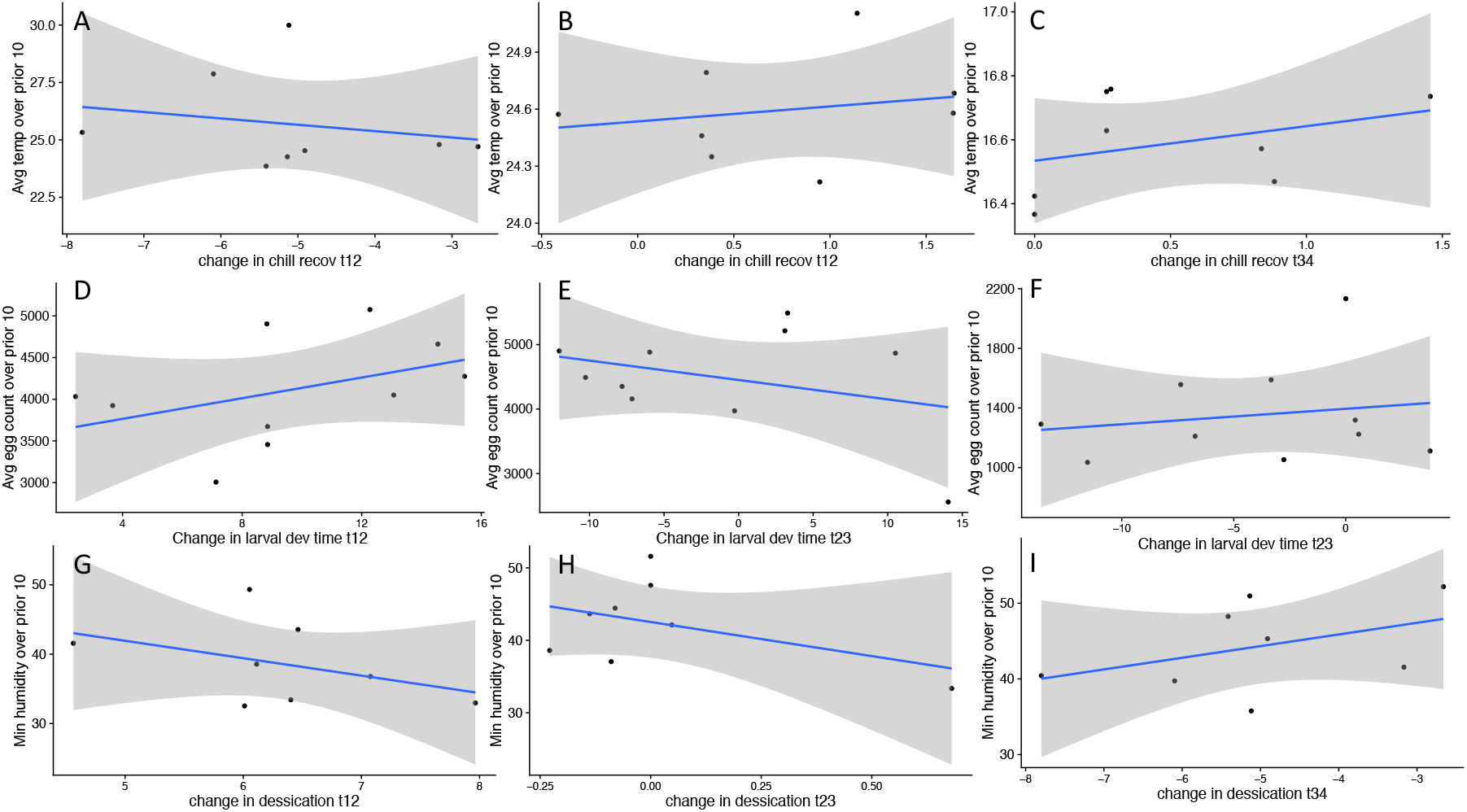
We assessed associations between cage to cage variation in environmental parameters and the pace of phenotypic evolution. Panels A, B, and C show the relationship between the average temperature in each cage (measured in 8 cages) over the 10 days preceding phenotyping and the change in genetic chill coma recovery time. Panels D, E, and F show the correlation between the average egg count and the change in genetic larval development rate. Panels G, H, and I show the correlation between the minimum humidity (measured in 8 cages) over the 10 days preceding phenotyping and the change in genetic desiccation tolerance. Overall, these associations did not uncover clear evidence of a specific environmental factor that drove cage to cage variation in evolutionary trajectories, suggesting that the agent of selection was something that did not vary strongly across cages, was not measured, or was shaped by several environmental factors over each time interval.

**Fig. S5.**
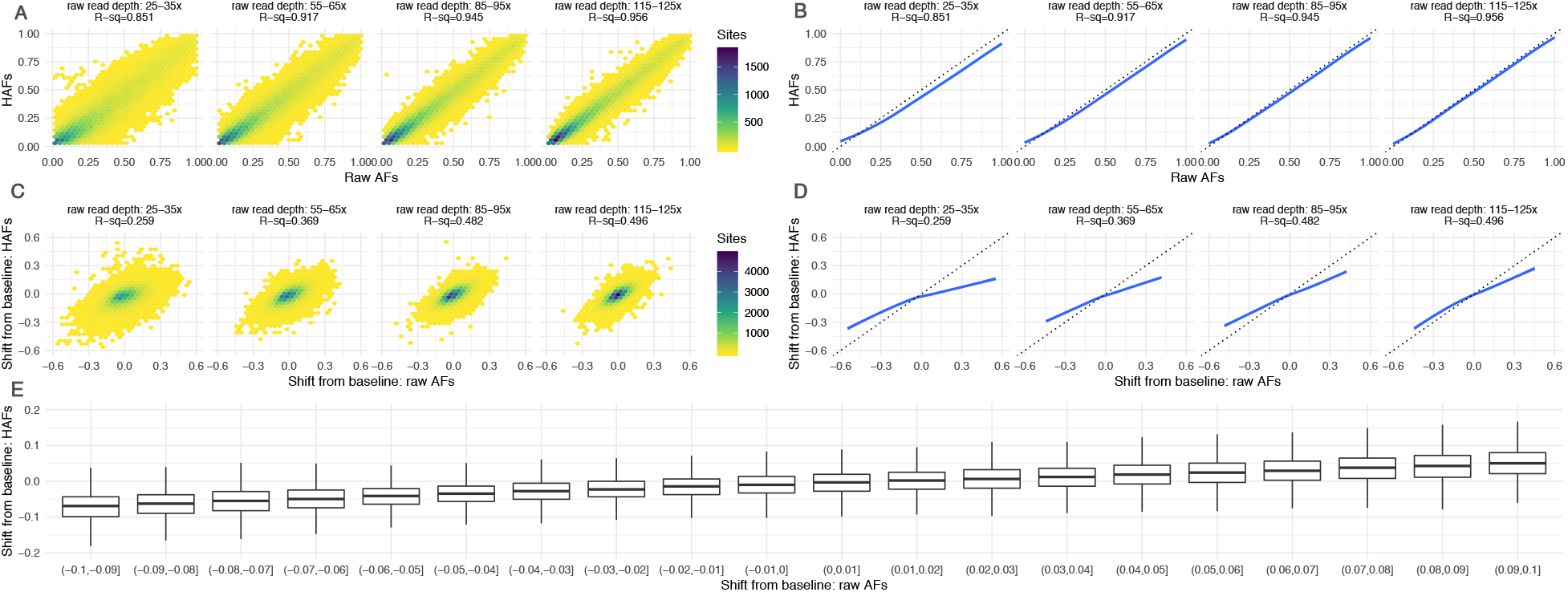
Haplotype-derived allele frequencies (HAFs; y-axis) obtained via low-coverage (~5x) sequencing of timepoint-5 samples followed by inference from founder haplotypes were compared to raw allele frequencies (x-axis) from deep re-sequencing of the same samples. Sites were binned by read depth in the deeply sequenced samples (separate panels). In all panels, concordance between HAFs and raw AFs increases as read depth of raw AFs increase, suggesting HAFs are effectively as accurate as raw AFs at >100x. A) Heatmaps of HAFs vs raw AFs for the same sample and site. B) Line of best fit (blue) for correlation between HAFs and raw AFs compared to line of perfect correlation (gray). C) Heatmaps of the shift between baseline and timepoint-5 calculated via HAFs vs raw AFs for the same sample and site D) Line of best fit (blue) for correlation between shifts from baseline calculated from HAFs vs raw AFs compared to line of perfect correlation (gray). E) Boxplots of HAF shifts binned by raw AF shift, at sites with raw read depth >=115x. All boxplots represent distributions with significantly different means (t-test p-values<.05).

**Fig. S6.**
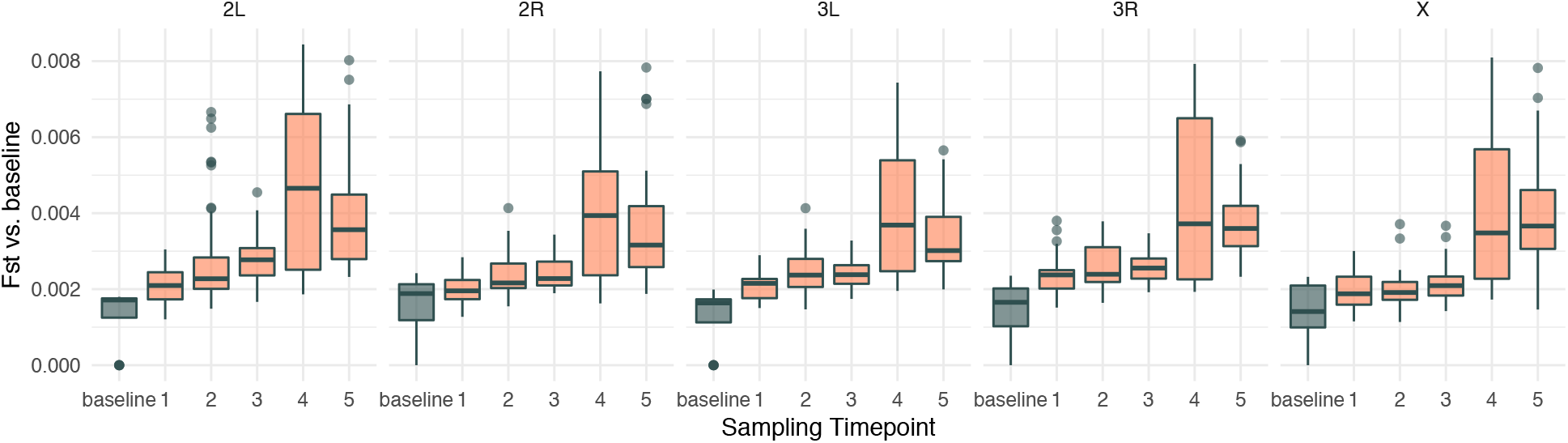
Distributions of chromosome-wide mean Fst between biological replicates from the baseline population (gray) or between experimental samples from each sampling timepoint and baseline samples (coral).

**Figure S7.**
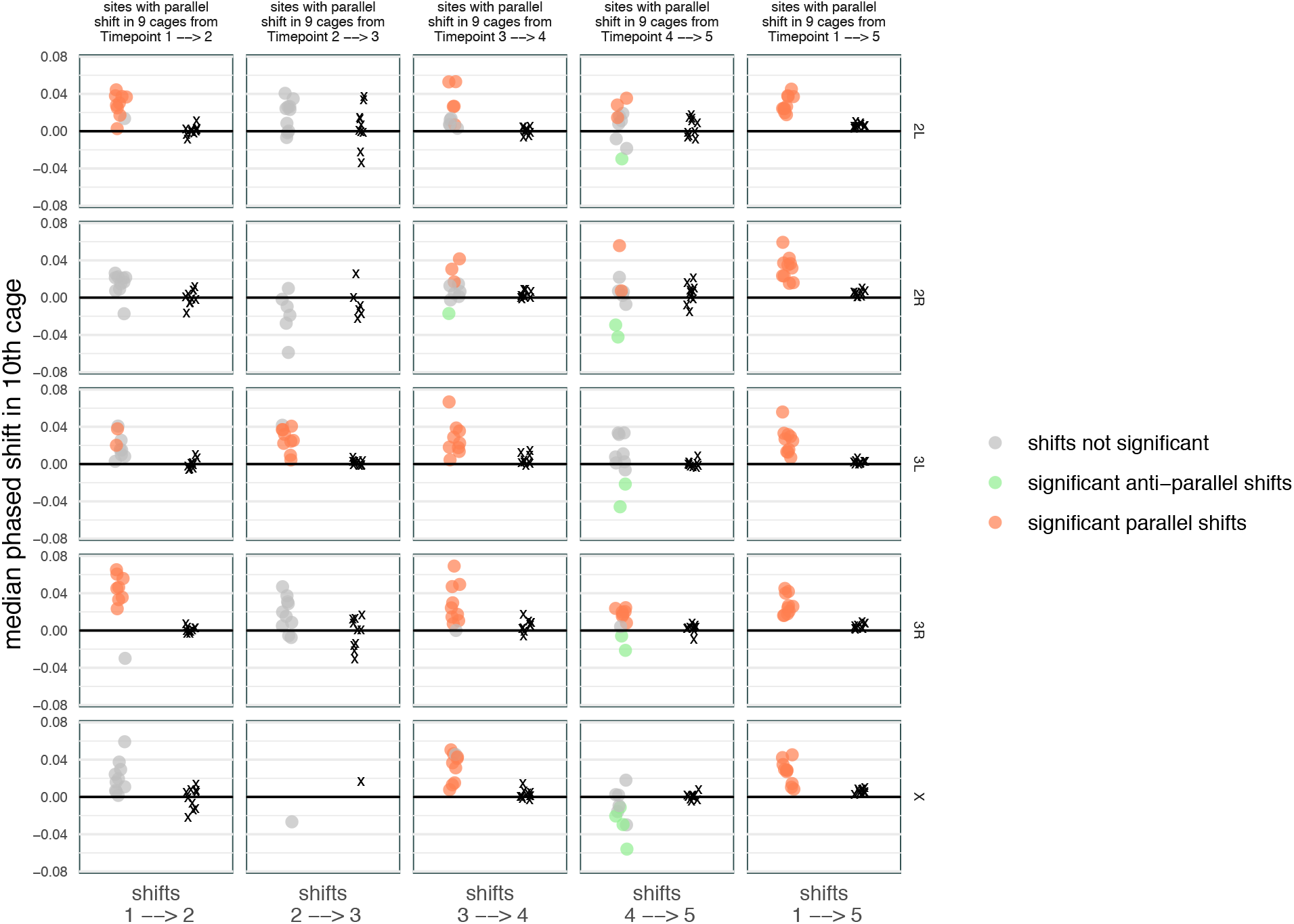
Per-chromosome leave-one out 10-fold cross-validation of significant sites. In each round, significantly parallel sites at each time-segment were identified using 9 of the 10 cages, then the shift at those sites in the 10th left-out cage was measured at the same time segment. In each case, the set of shifts at parallel sites was compared to shifts at control sites matched for chromosome and initial frequency to determine whether shifts in the left-out cage at parallel sites were also significantly parallel (orange) or significantly anti-parallel (green). Median shift for each set of parallel sites (circles) and control sites (x marks) on each chromosome (rows) at each time segment (columns) are plotted for each left-out cage.

**Fig S8.**
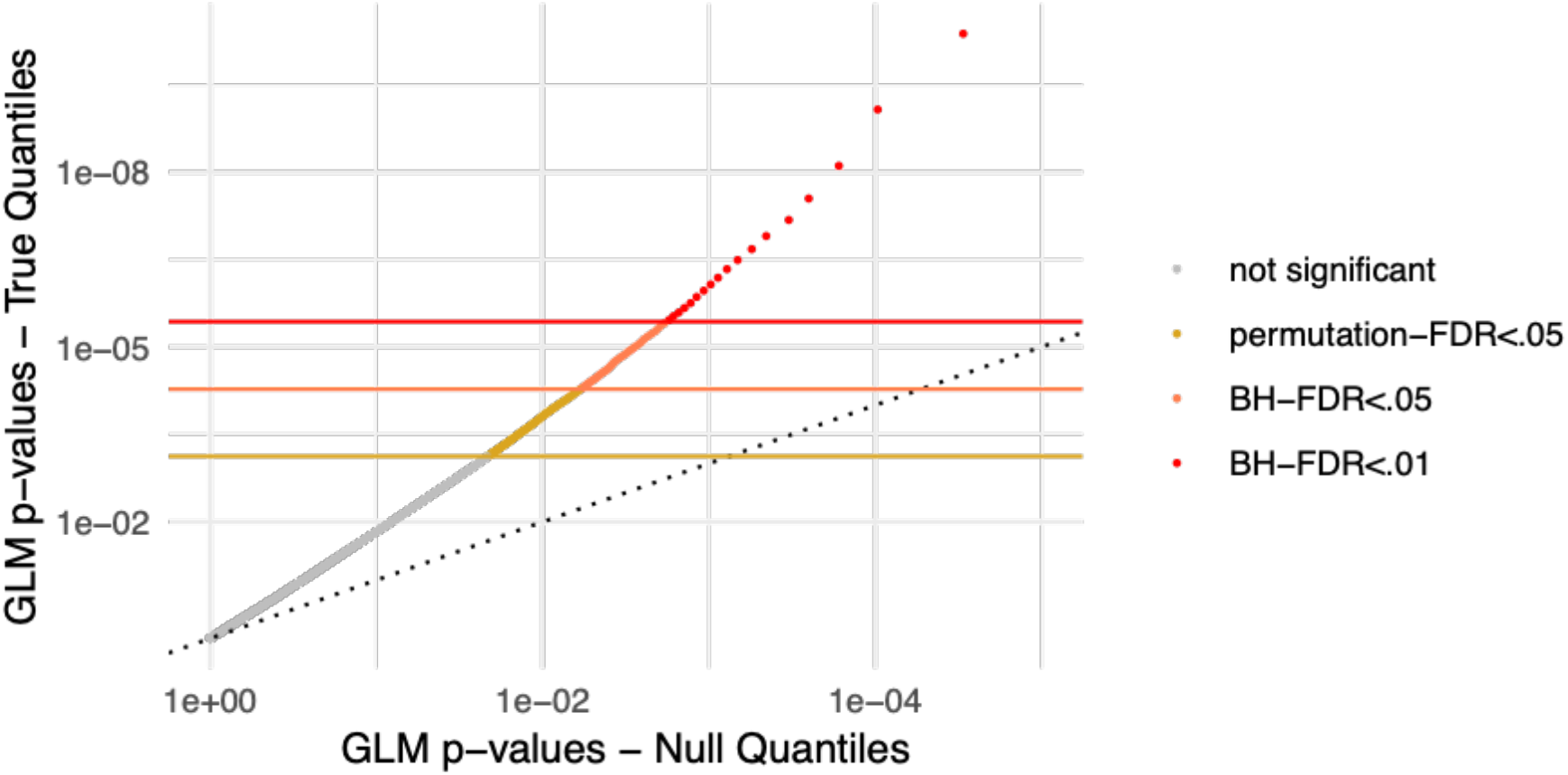
QQ plots for per-SNP GLM p-values giving the significance of a parallel shift across 10 replicate cages for true data (y-axis) and null model (x-axis) where timepoint labels for each site were shuffled across samples before fitting the GLM. Color of each point indicates whether the p-value for the true quantile passes various FDR thresholds.

**Figure S9.**
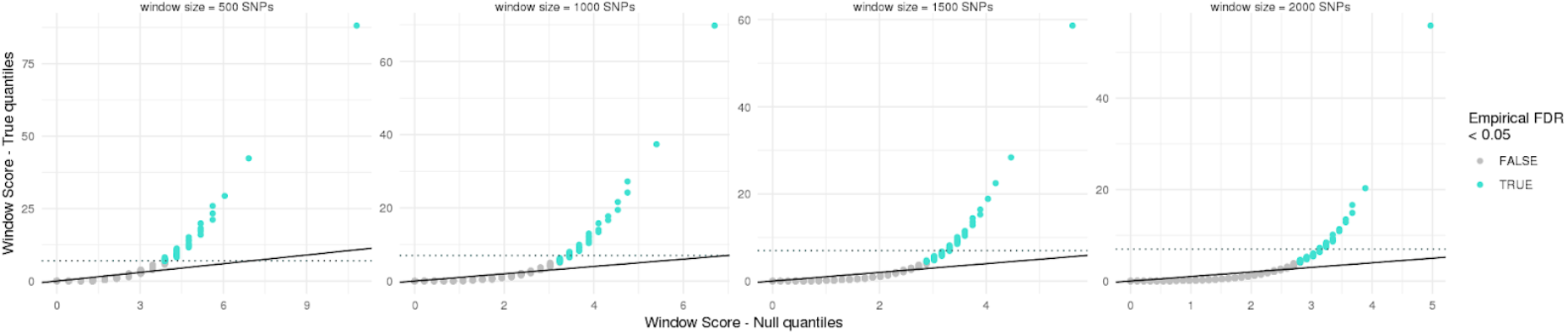
QQ plots of the distribution of significant sites in various equi-SNP sized sliding windows. Each SNP was scored (0, 1, 2, or 3) according to significance of parallelism at each time segment (see Methods). SNP-scores were averaged across consecutive SNPs to generate a window score. True window score quantiles (y-axis) were compared to quantiles from a null distribution generated by randomly shuffling SNP labels.

**Figure S10.**
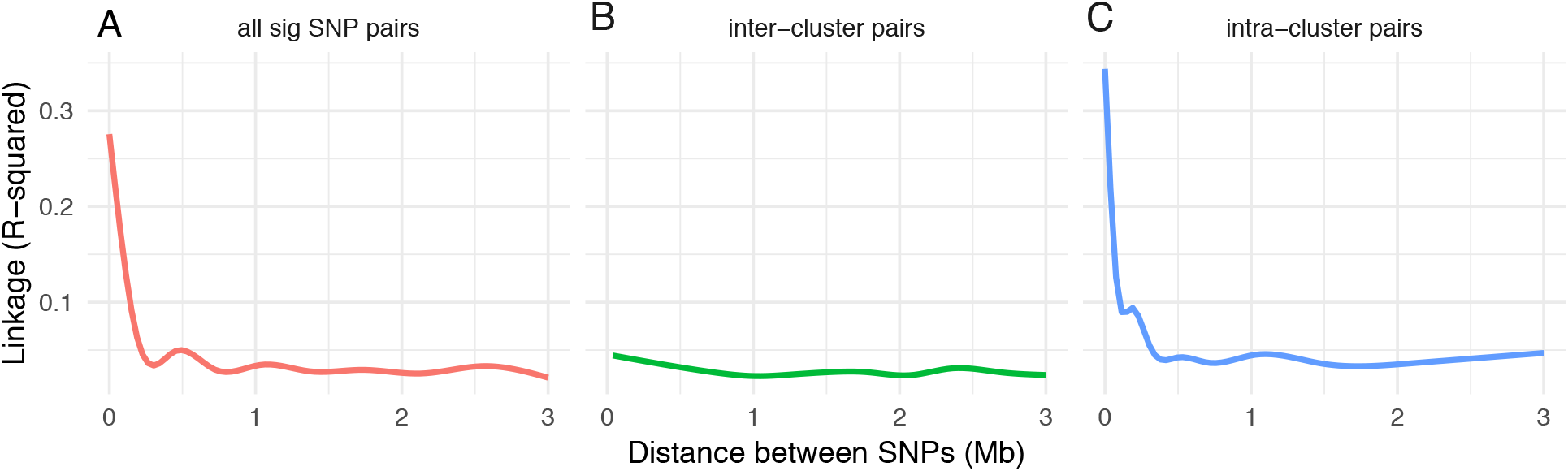
Smoothed average of linkage values as a function of distance between SNPs, measured between **A)** all pairs of significant (FDR<.05) SNPs on the same chromosome identified at the same time segment, **B)** pairs of SNPs on the same chromosome identified at the same time segment that were assigned to different clusters, and **C)** pairs of SNPs on the same chromosome identified at the same time segment that were assigned to the same cluster.

**Figure S11.**
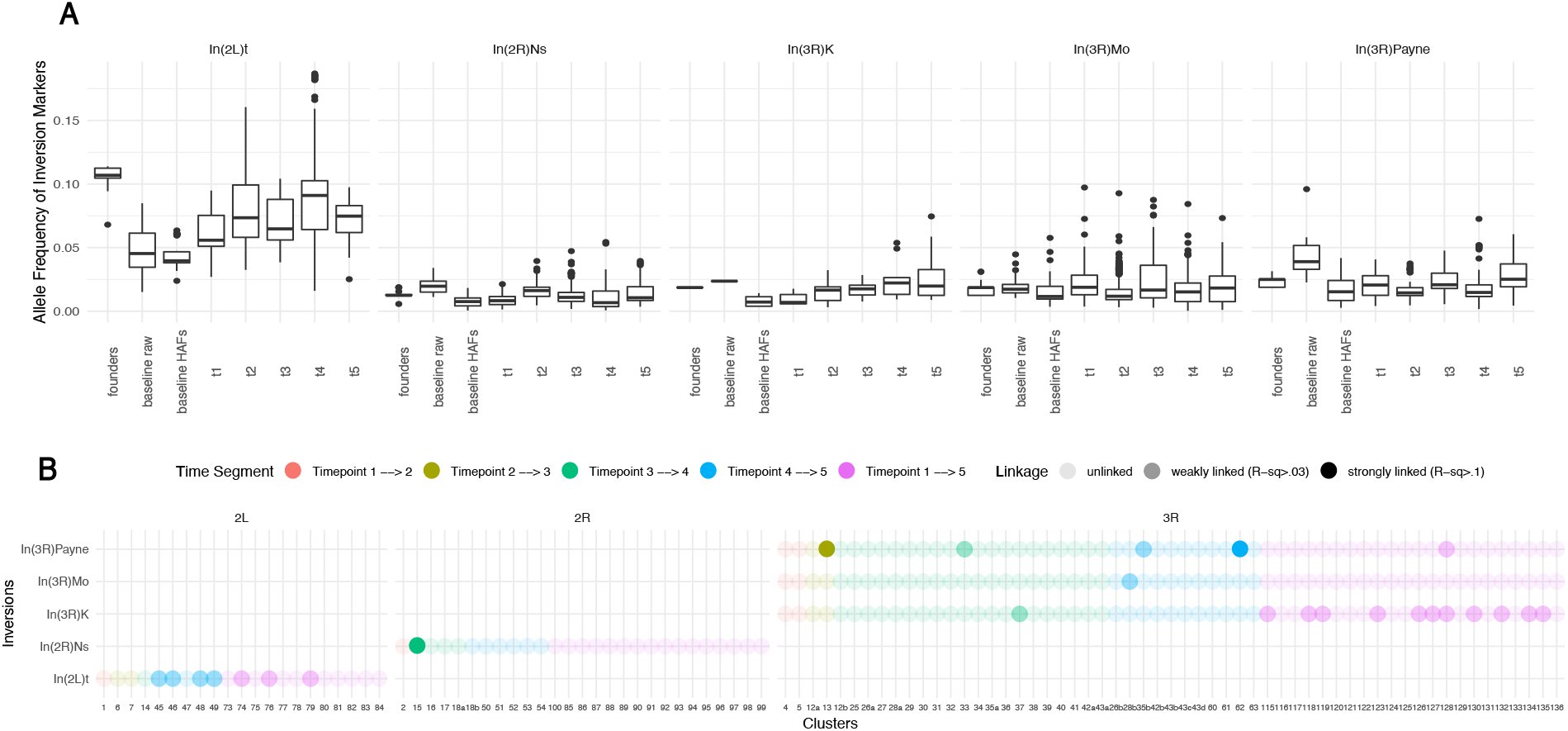
**A**) Distribution of the allele frequencies of inversion markers in the founder lines, baseline population, and across cages at each timepoint. **B**) Linkage between clusters (x-axis) and inversions (y-axis). Dots are colored by time segment of cluster identification and shading indicates whether clusters are unlinked, weakly linked (average R-squared between significant parallel SNPs and inversion markers is > 0.03) or strongly linked (average R-squared > 0.1).

**Figure S12.**
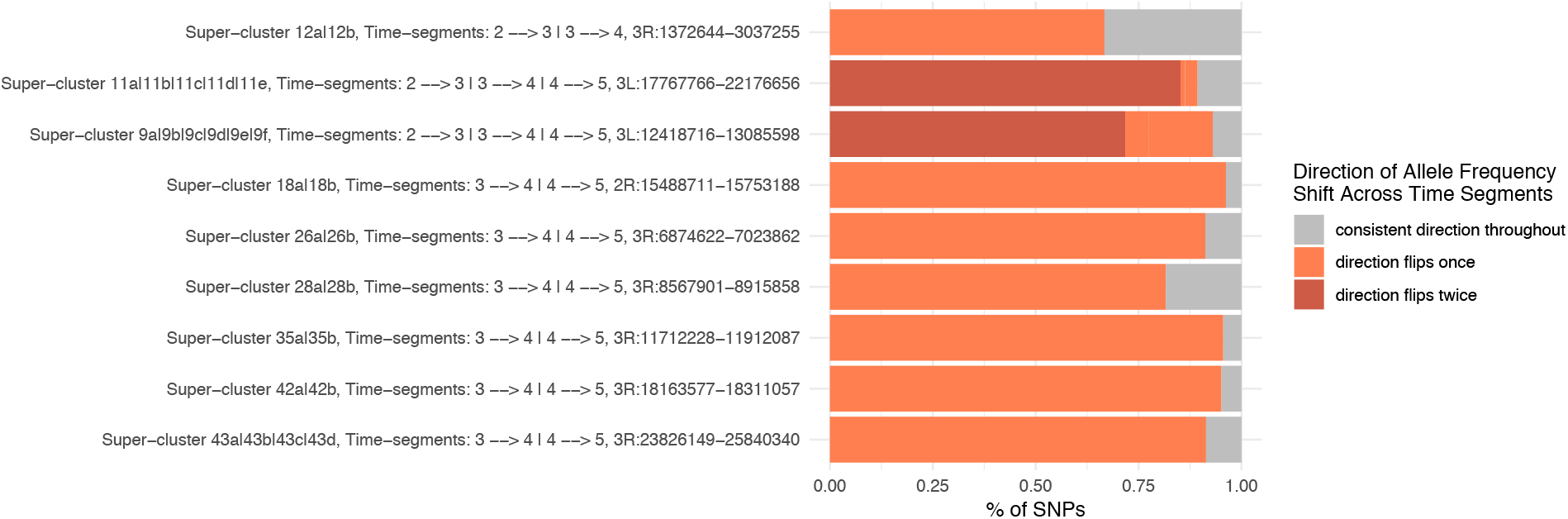
Proportion of SNPs in the intersection of linked clusters from different time segments (aka superclusters) that continue shifting in the same direction across months (gray) or flip direction (orange). Two superclusters involve linked clusters from three different time segments (T_2_→T_3_, T_3_→T_4_, and T_4_→T_5_); for these superclusters, color indicates the consistency of direction between T_2_➔ T_3_ and T_3_→T_4_, followed by the consistency of direction between T_3_→T_4_ and T_4_→ T_5_ (i.e., flips, same).

**Figure S13.**
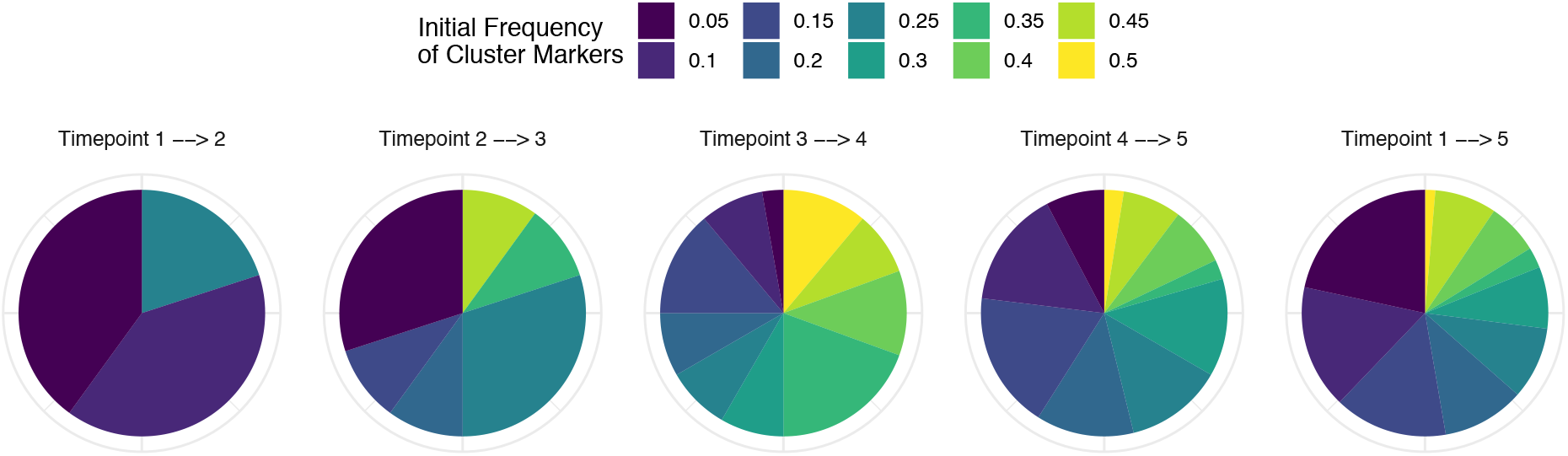
Distribution of the initial minor allele frequencies of marker SNPs for clusters identified at each time segment.

